# Comparative basolateral amygdala connectomics reveals dissociable single-neuron projection patterns to frontal cortex in macaques and mice

**DOI:** 10.1101/2023.12.18.571711

**Authors:** Zachary R Zeisler, Kelsey A Heslin, Frederic M Stoll, Patrick R Hof, Roger L Clem, Peter H Rudebeck

## Abstract

The basolateral amygdala (BLA) projects to the frontal cortex (FC) in both rodents and primates, but the comparative organization of single-neuron BLA-FC projections is unknown. Using a barcoded connectomic approach, we found that BLA neurons are more likely to project to multiple distinct parts of FC in mice than in macaques. Further, while single BLA neuron projections to nucleus accumbens are similarly organized in mice and macaques, BLA-FC connections differ.

## Main text

Basolateral amygdala (BLA) is a key hub for affect in the brain^1–3^ and dysfunction within this area contributes to a host of psychiatric disorders^4,5^. BLA is extensively interconnected with frontal cortex^6–9^, and some aspects of its function are evolutionarily conserved across rodents, anthropoid primates, and humans^10^. Neuron density in BLA is substantially lower in primates compared to murine rodents^11^, and frontal cortex is dramatically expanded in primates, particularly the more anterior granular and dysgranular areas^12–14^. Yet how these anatomical differences influence the projection patterns of single BLA neurons to frontal cortex across rodents and primates remains unknown. Establishing the shared connectivity patterns across species is essential for interpreting experimental findings from rodents to monkeys and *vice-versa*.

To characterize these projections, we conducted parallel MAPseq experiments in laboratory mice and rhesus macaques by making injections of barcoded Sindbis virus into BLA (**Figure 1a**). MAPseq^15^ uses a Sindbis virus vector to infect neurons with unique RNA barcode sequences to profile single-neuron projection patterns at scale, enabling detection of axonal branching and sufficient throughput to characterize thousands of projections in a small number of animals^15–18^. Following perfusion, the brains were then extracted, blocked, and sectioned in the coronal plane^15,16^. Areas of interest in amygdala, frontal cortex, striatum, thalamus, and hippocampus were then dissected (**Tables 1 and 2**). We specifically obtained samples from cortical and subcortical areas that are thought to be analogous or possibly homologous across mice and macaques to enable direct species comparison^12,19,20^. RNA was extracted from dissected samples and sequenced. From this, a matrix of RNA barcode counts across the samples was constructed, a threshold applied, and all data were collapsed across subjects within a species (see **Online Methods**). Before conducting further analyses we confirmed that: 1) few barcodes were recovered from samples taken from cerebellum in macaques and primary visual cortex (V1) in mice, our two control areas (**Supplemental Figure 1A**), 2) thresholds were appropriately applied (**Supplemental Figure 1B)**, and 3) the pattern of barcode expression across samples was primarily unique indicating that single virus particles likely infected individual neurons (**Supplemental Figure 1C**). Overall, these analyses confirmed that MAPseq was working similarly and as effectively in both mouse and macaque BLA.

**Figure 1:**
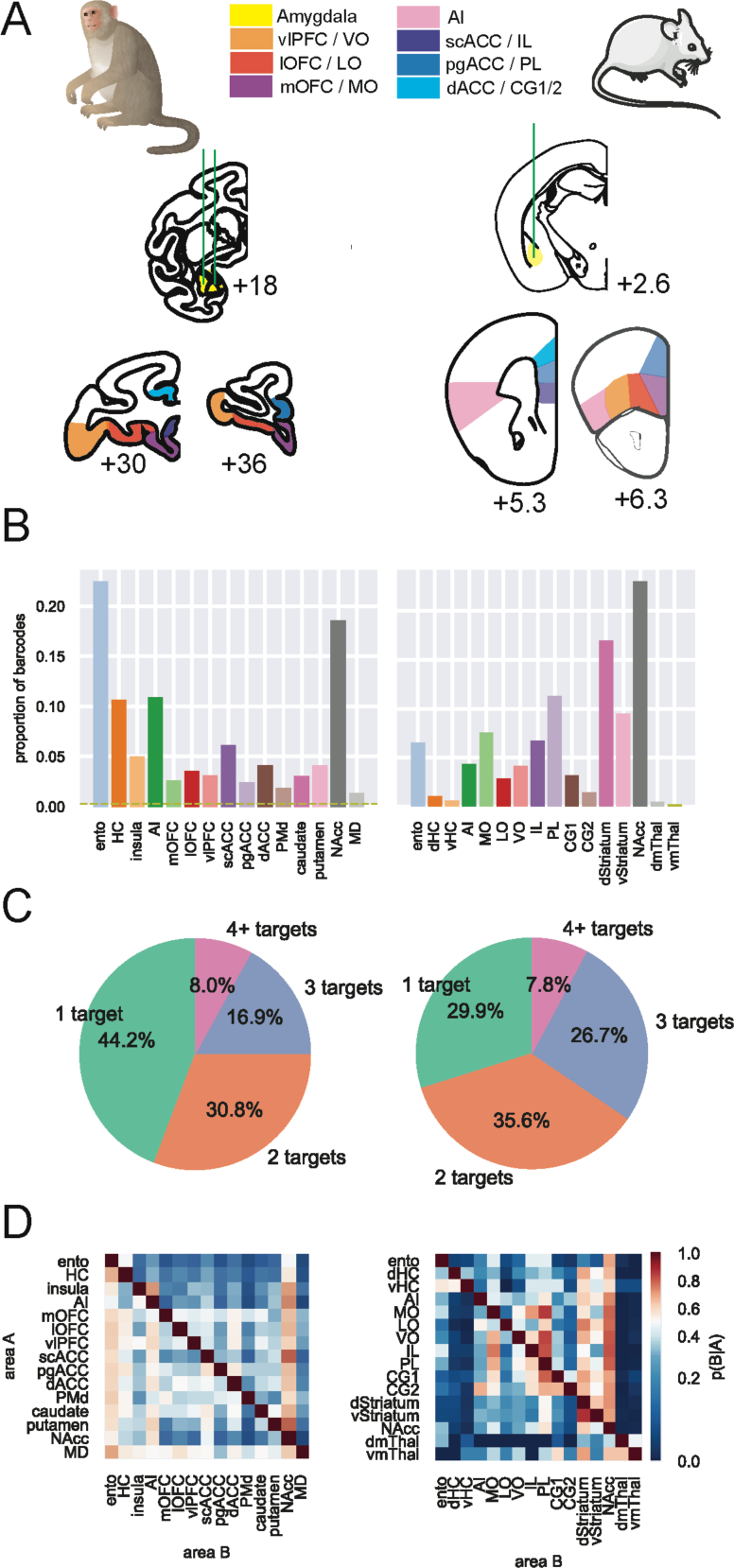
Single BLA neurons projections to frontal cortex in mice and macaques. A) Schematic of injection (middle) and target (bottom) sites in macaques (left) and mice (right). Green vertical lines indicate injection tracks, and interaural distance is provided in mm. Samples were collected from amygdala (yellow), vlPFC/VO (orange), lOFC/LO (red), mOFC/MO (purple), AI (pink), scACC/IL (dark blue), pgACC/PL (blue), and dACC/CG1&2 (light blue). B) Overall projection strength as measured by proportion of barcodes transported to each target area. C) Number of targets for each neuron. 1-target neurons were found in amygdala and one other brain area, 2-targets in amygdala and two other brain areas, and so on. Mouse BLA neurons were significantly more likely to have multiple targets than macaques (Fisher’s exact test: *p* < 0.001). D) Conditional probabilities of branching neurons. Probability that a barcode found in Area A (vertical axis) is also found in Area B (horizontal axis) is indicated by color, such that warmer colors indicate more frequent overlap.

**Table 1.** Definitions of brain regions collected from macaques. Here, we provide the names, abbreviations, and anatomical features used to determine the brain regions collected for the macaque MAPseq experiments. Repeated from Zeisler et al., 2023.^16^.

| Brain Region | Abbreviation | Approximate Walker's Areas | A/P boundaries | M/L boundaries |
| --- | --- | --- | --- | --- |
| Amygdala | n/a |  | All nuclei across entire A/P extent |  |
| Entorhinal cortex | ento |  | From the anterior tip to a point adjacent to the mid-hippocampus. | Midline to fundus of rhinal sulcus |
| Hippocampus | HC |  | Anterior tip to the middle of hippocampus |  |
| Medial orbitofrontal cortex | mOFC | 11m, 13a/b, 14 | From emergence of medial orbital sulcus to its disappearance | Between rostral sulcus and medial orbital sulcus |
| Lateral orbito-frontal cortex | IOFC | 11l, 12m/r, 13m/l | From emergence of lateral orbital sulcus to its disappearance | Between fundus of medial sulcus and fundus of lateral orbital sulcus |
| Ventrolateral prefrontal cortex | vIPFC | 45, 12o, 12l | From emergence of lateral orbital sulcus to its disappearance | Between lateral orbital sulcus and lateral convexity of ventral surface |
| Dorsal anterior cingulate cortex | dACC | 24 | From emergence of cingulate sulcus until area 25 disappears from medial wall/emergence of septum. | Dorsal and ventral banks of cingulate sulcus, and cortex immediately dorsal to corpus callosum |
| Subcallosal anterior cingulate cortex | scACC | 25 |  | Medial surface, ventral to corpus callosum, medial/dorsal to medial orbital sulcus |
| Perigenual anterior cingulate cortex | pgACC | 32 | Anterior to corpus callosum | Cortex on medial surface of the frontal lobe between the rostral sulcus and the cingulate sulcus |
| Dorsal premotor cortex | PMd | 6 | Posterior to emergence of dorsal arcuate sulcus to emergence of arcuate spur | Dorsal convexity to arcuate spur |
| Caudate nucleus | caudate |  |  |  |
| Putamen | n/a |  |  |  |
| Nucleus accumbens | NAcc |  | From presence of internal capsule to the full separation of caudate nucleus and putamen | Ventromedial portion of striatum |
| Agranular insula | AI |  | Orbital surface after disappearance of medial and lateral orbital sulci | From medial corner of orbital surface to approximate location of fundus of lateral sulcus |
| Insula | n/a |  | Posterior to frontal/temporal junction to midpoint of hippocampus | Most medial vertically-oriented cortex in depth of lateral fissure |
| Medial dorsal thalamus | MD |  | From the optic tract joining the cerebral white matter to the lateral habenula | Dorsal and medial thalamic nuclei |

**Table 2.**
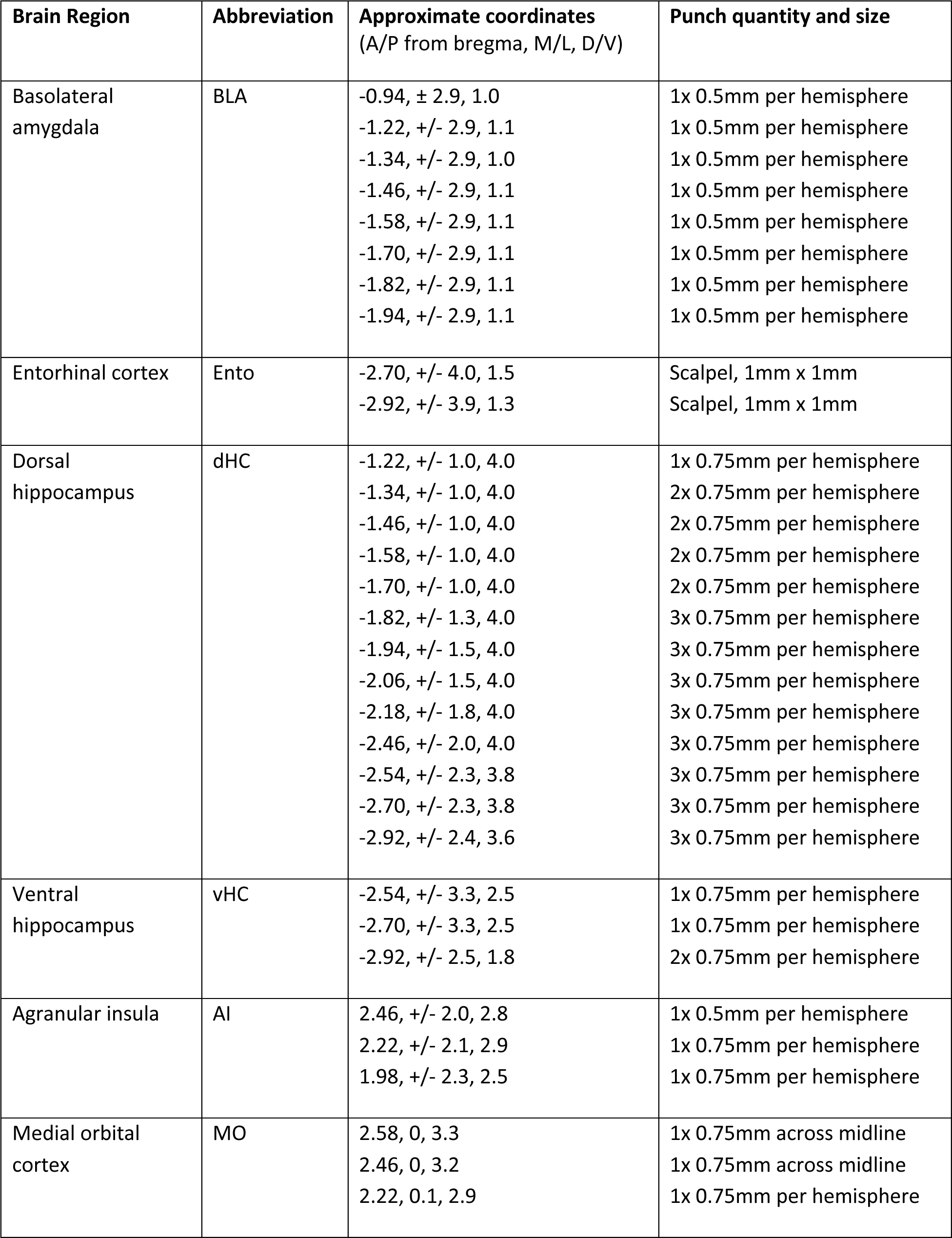
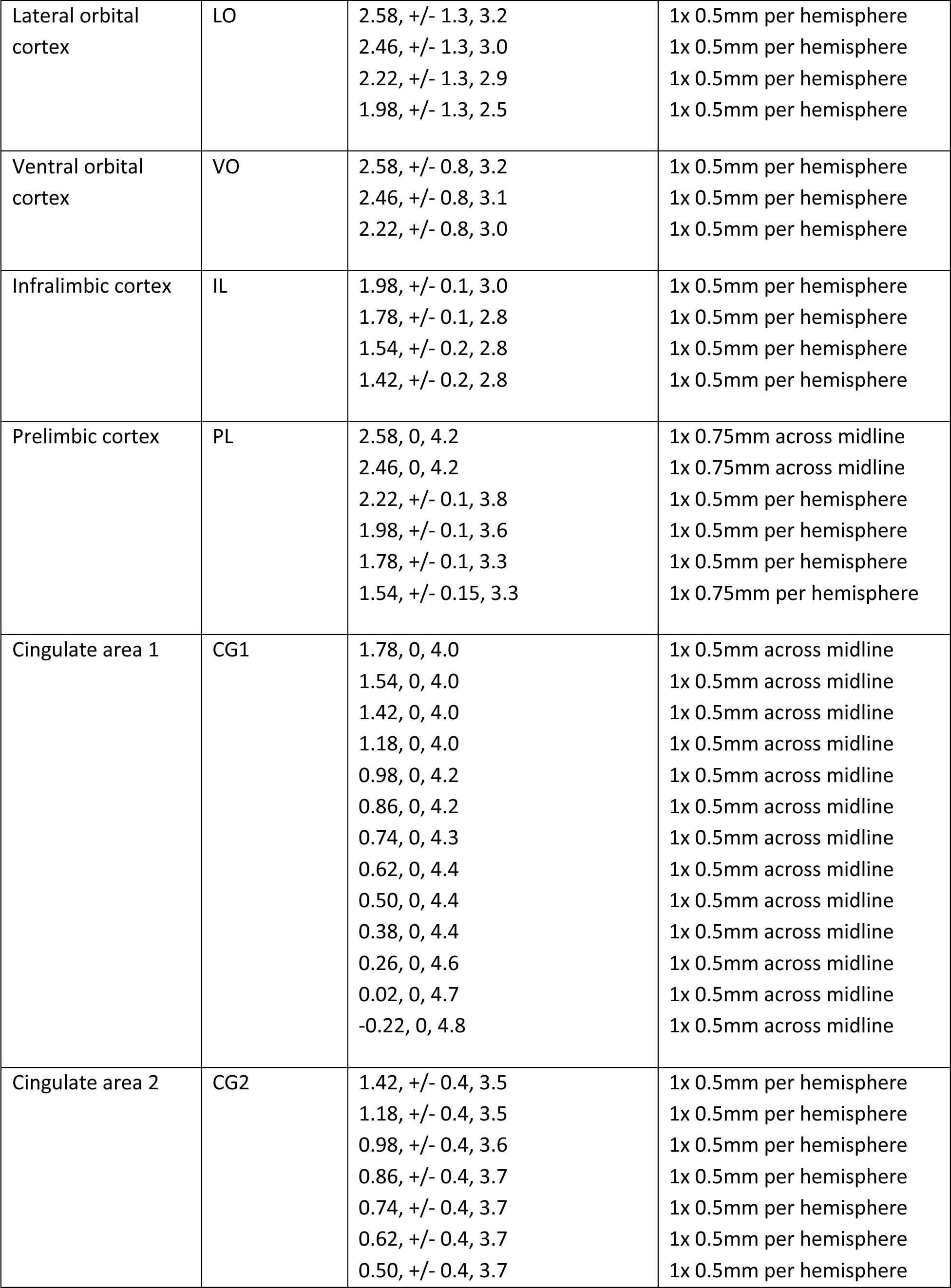
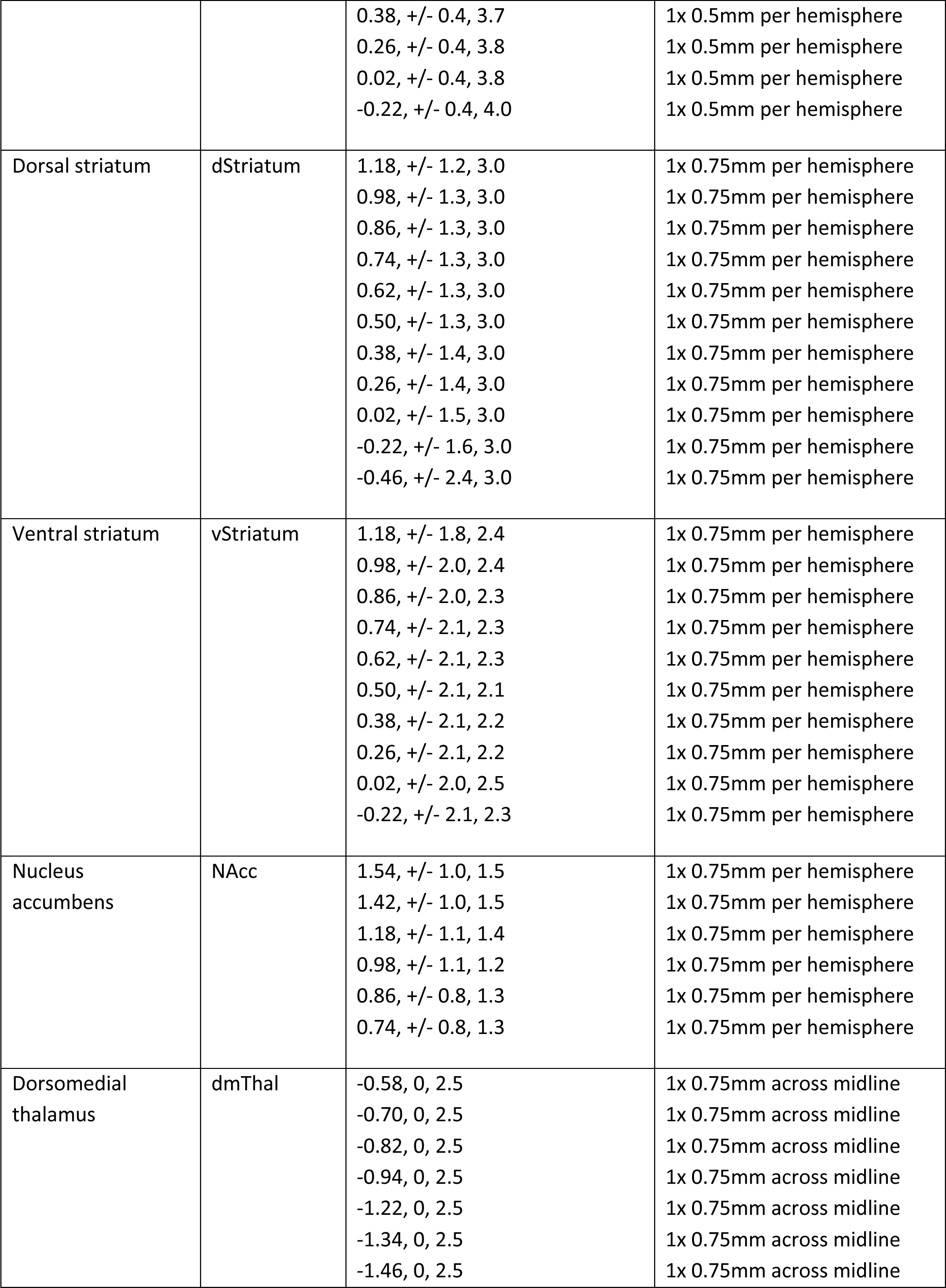
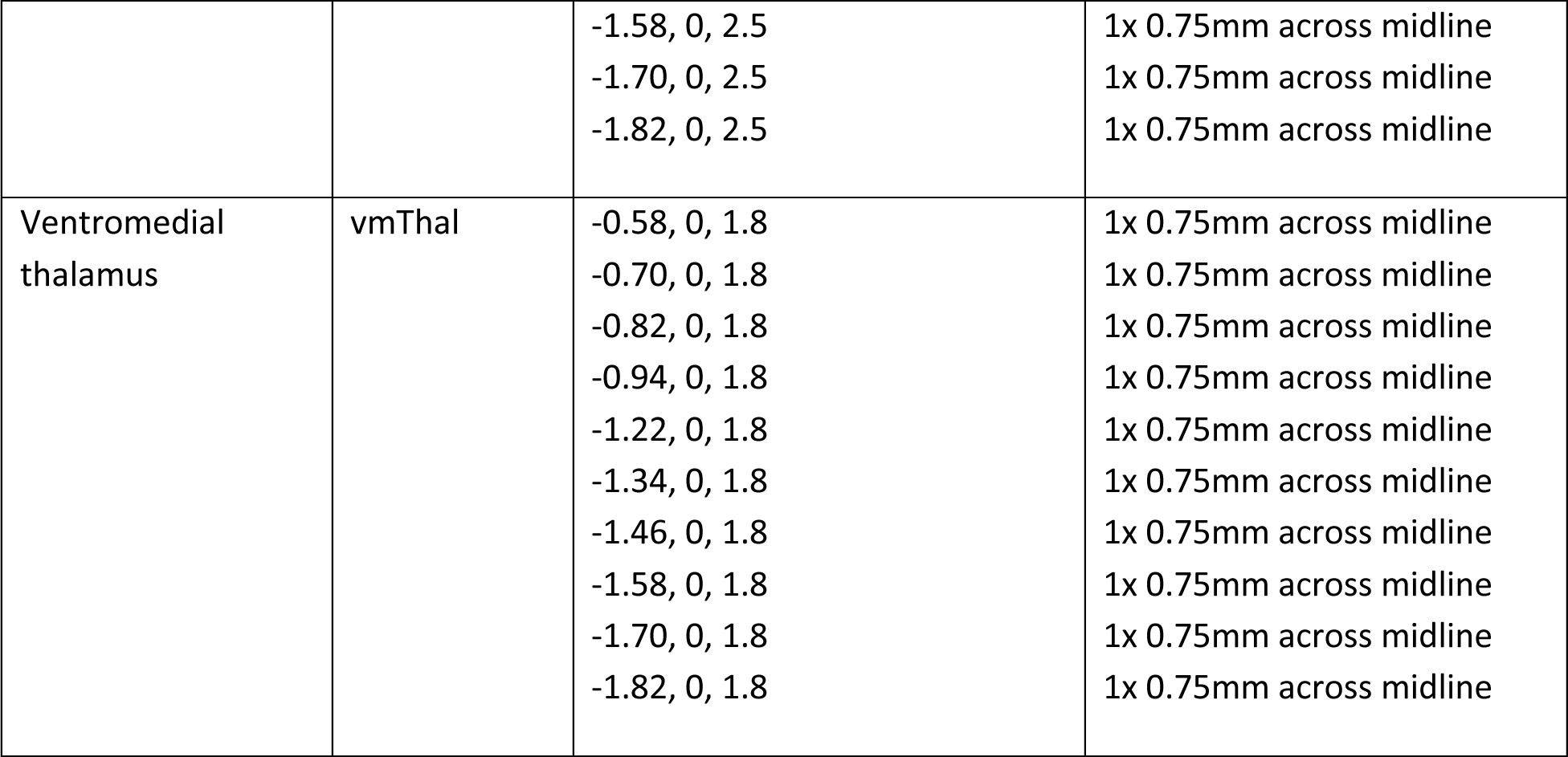
Definitions of brain regions collected from mice. Here, we provide names, abbreviations, approximate coordinates (from ref.^42^) of the center of the punches, and the number and size of punches collected for the mouse MAPseq experiments. Each set of coordinates refers to a punch taken from a coronal section, all of which from a given area were combined into a single sample for processing and sequencing.

A total of 1,674 and 3,115 unique barcodes were recovered from BLA across mice and macaques, respectively. In both species, the proportion of barcodes that we recovered from target locations in frontal cortex, striatum, temporal cortex, and thalamus closely aligned with prior tract tracing work on the projections of BLA^7^. The majority of recovered barcodes – which we interpret as projections from single BLA neurons - were found in striatum, including nucleus accumbens (NAcc), compared to frontal cortical areas or thalamus (**Figure 1B**). In addition to these bulk tracing patterns, we also found that more mouse BLA neurons branched to multiple locations than in macaques (**Figure 1C**). Specifically, although all sequenced neurons were more likely to branch to more than one target, BLA neurons in mice were more likely to project to more than two or three targets compared to macaques. Thus, the larger frontal cortex in macaques is not associated with a higher degree of collateralization.

To further explore the organization of the connections of single BLA neurons to multiple areas in mice and macaques, we next computed the likelihood that a barcode found in one target structure would also be found in another (**Figure 1D**). Here, we found similarities in the projection patterns of single BLA neurons to subcortical structures between the two species. Notably, neurons that projected to NAcc were highly likely to also project to all parts of frontal cortex in both mice and macaques. Despite this similarity, there were species distinctions in how BLA neurons projected to frontal cortex. For instance, BLA neurons projecting to prelimbic cortex (PL) in mice were highly likely to also project to other parts of frontal cortex (**Figure 1D**). BLA projections to the area in macaques thought to be homologous to PL^19^, perigenual anterior cingulate cortex (pgACC, Walker’s area 32) were by contrast very unlikely to project to other parts of frontal cortex. We also found a clear difference between the caudate in macaques and the dorsal striatum in mice. Specifically, dorsal striatum in mice was far more likely to receive BLA input and share that input with other frontal areas. By comparison, macaque caudate nucleus was only weakly innervated by BLA and received largely input unlikely to be shared with other areas, potentially highlighting the need for careful comparison of findings from this region across species.

Next, focusing on the patterns of projections from BLA to frontal cortex, we looked at how single BLA neurons separately connected to medial and ventral frontal cortex (see **Online Methods** and Tables for definitions of these regions, illustrated in **Figure 2** and **Supplemental Figure 2**, respectively). BLA-medial frontal circuits (**Figure 2A**) in rodents and primates are involved in aspects of social behavior^21–24^ and defensive threat conditioning^25,26^. Notably, cross-species comparisons indicate that the proposed homologues PL/pgACC and infralimbic cortex (IL)/subcallosal ACC (scACC; Walker’s area 25) differentially contribute to defensive threat conditioning across species, where the specific functions subserved by these areas is reversed between rodents and primates^19^. Mirroring that functional distinction, we found that BLA projections to these parts of medial frontal cortex exhibited a reversal in the pattern of specific or branching connections. While scACC received the most specific BLA input with respect to the other medial frontal areas in the macaque (**Figure 2B**), it was the mouse PL which received a similar proportion of input not shared with the other medial areas.

**Figure 2:**
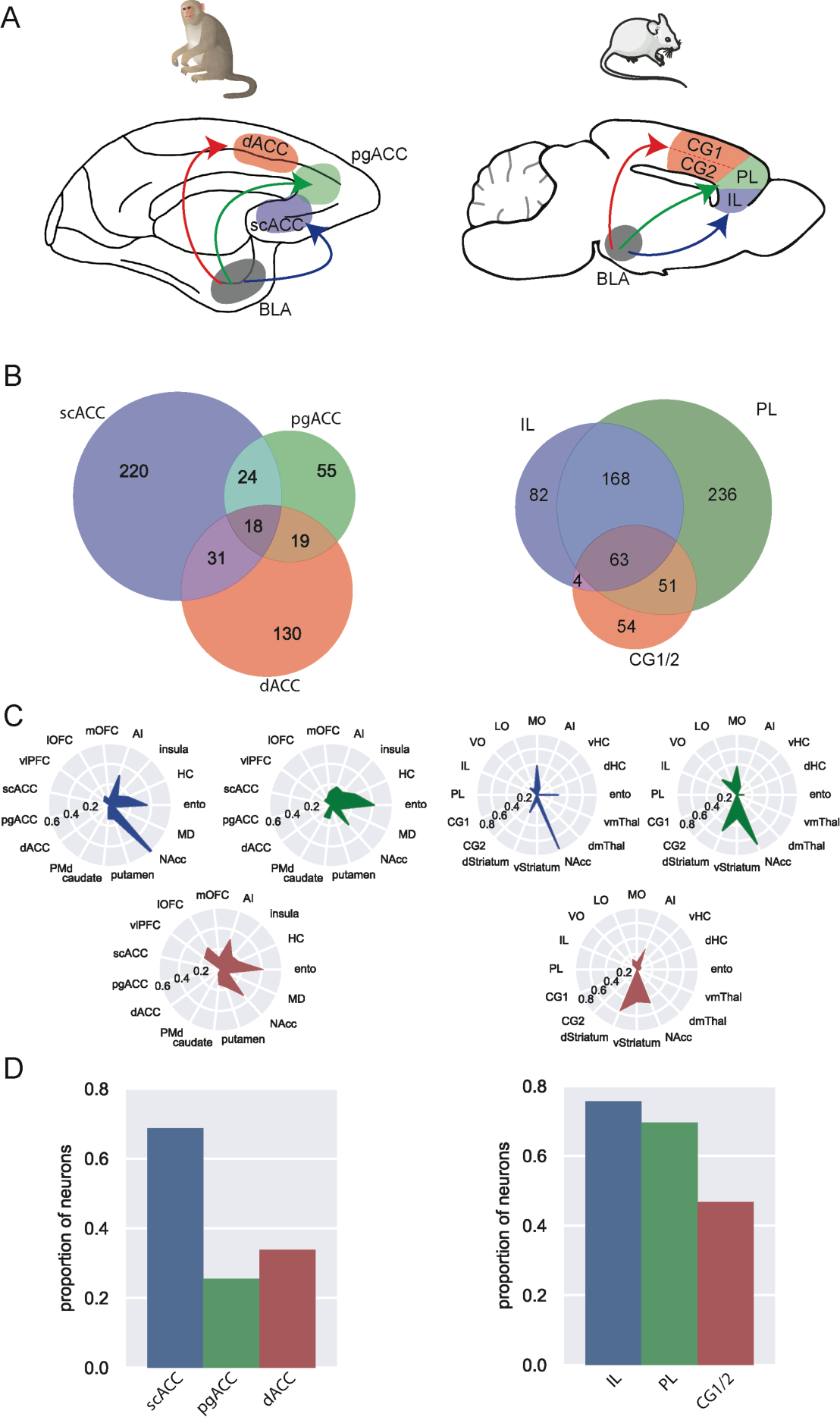
The organization of single BLA neuron projections to medial frontal cortex in mice and macaques. A) Schematic of populations of medial-projecting neurons in macaques (left) and mice (right). B) Venn diagrams of within-medial frontal cortex branching. scACC was least likely to branch within medial frontal cortex in macaques (z-test for proportions, z = 6.38, *p* < 0.0001), while PL in the mouse instead received the most specific input (z = 4.64, *p* < 0.0001). C) Likelihood of medial frontal-projecting neurons branching to project to other areas. This plot excludes those neurons which branch between multiple medial frontal areas, such that the populations are non-overlapping. dACC-projecting neurons were most likely to branch to lOFC (dACC vs pgACC: z = 2.72, *p* = 0.0097; dACC vs scACC: z = 3.80, *p* < 0.001) and vlPFC (vlPFC; dACC vs pgACC: z = 2.04, *p* = 0.061; dACC vs scACC: z = 3.52, *p* < 0.001); whereas PL-projecting neurons were most likely to branch to VO (PL vs CG1/2: z = 3.22, *p* = 0.0013; PL vs IL: z = 2.82, *p* = 0.0048). D) Likelihood of ventral frontal-projecting neurons projecting to NAcc. scACC-projecting neurons were most likely to also project to NAcc (scACC vs dACC: z = 2.03, *p* = 0.005), whereas both IL- and PL-projecting neurons were equally likely to branch to NAcc (PL or IL vs CG1/2: z = 3.33, *p* < 0.001).

Similarly, when we focused on the non-overlapping populations of neurons projecting to each medial area and assessed their connectional fingerprints (**Figure 2C**), we identified many differences across motifs. Of particular interest to us were motifs which included subdivisions of the orbitofrontal cortex (OFC) which are well-documented to be involved in value-based decision-making in both rodents and macaques^27^. We found that of the BLA neurons projecting to macaque medial frontal cortex, those to dorsal ACC (dACC) had the highest likelihood of also projecting to lateral OFC (lOFC) and ventrolateral prefrontal cortex (vlPFC). In mice, the combinations differed. Instead, BLA neurons projecting to PL were more likely to also project to ventral orbital cortex (VO) rather than the area thought to be the homologues of macaque dACC, (i.e. areas CG1 and CG2).

Finally, we focused on BLA-medial frontal projections to NAcc, a highly-conserved subregion of the striatum^28^ which has been implicated in substance use behaviors^29–31^ and learning from salient experiences^32,33^ across species. In macaques, we found that BLA neurons projecting to scACC (area 25; the IL homologue) are highly likely to also innervate NAcc (**Figure 2D**), while neurons projecting to pgACC (area 32; the PL homologue) were not. In mice, however, both IL- and PL-projecting neurons were equally likely to branch to NAcc. Here, much like in the overall branching proportions (**Figure 1C**), we observe that mouse BLA neurons are more likely to project widely while macaque BLA neurons have more specific projection patterns. Taken together, these results highlight distinctions in the organization of BLA projections to medial frontal cortex and that such differences appear to align with cross-species functional studies, in which primate scACC and pgACC play different roles in regulating affect than their rodent counterparts, IL and PL^19^.

To determine whether these distinctions were unique to medial frontal cortex, we performed the same analyses on BLA projections to ventral frontal cortex (see **Online Methods**). While interaction between BLA and ventral frontal cortex is essential for goal-directed behaviors in both species^34,35^, we discovered clear distinctions in BLA-ventral frontal projections (**Supplemental Figure 2A**). In macaque ventral frontal cortex, we found a gradient across areas where the majority of neurons projecting to agranular insula cortex (AI) only targeted this area, whereas BLA neurons targeting ventrolateral prefrontal cortex (vlPFC) were highly likely to branch to other parts of ventral frontal cortex (**Supplemental Figure 2B**, left). The same pattern was largely absent in mice (**Supplemental Figure 2B**, right). Specifically, MO received the highest proportion of input not shared with the rest of ventral frontal cortex, whereas BLA neurons targeting VO and LO were most likely to branch. That vlPFC and VO both receive BLA input frequently shared among other ventral frontal areas is consistent with their shared functions in adapting choice behavior in probabilistic environments^36,37^. When we compared the connectional fingerprints of mouse and macaque ventral frontal projecting single BLA neurons (**Supplemental Figure 2C**), we again found striking differences. For instance, BLA neurons projecting to mouse area LO, which is thought to be analogous to the caudal part of macaque lOFC^38^ were highly likely to project to PL as well as IL. Such a pattern was almost completely absent in macaques. Instead, BLA neurons projecting to lOFC were more likely to target either dACC (area 24) or scACC (area 25) and not pgACC (area 32), the homologue of PL^12^. Across BLA neurons projecting to ventral frontal cortex, we also observed cross-species distinctions in the strength of projections to NAcc (**Supplemental Figure 2D**).

Thus, despite substantial functional similarities between BLA-ventral frontal circuits, there were significant distinctions in the patterns of BLA projections to these areas across species. Overall, our findings reveal that single BLA neurons in macaques are less likely to collateralize to multiple areas of frontal cortex compared to mice (**Figure 1C**). Such a difference is important as it indicates that BLA-frontal circuits in macaques are not merely a scaled-up version of those present in mice. Instead, the connections emanating from BLA to the expanded frontal cortex are more segregated in macaques (**Figure 2** and **Supplemental Figure 2**). One potential explanation for this difference is that frontal areas in macaques^12^ have more specialized functions and might require more specific inputs. Our results, thus, provide anatomical evidence for how findings related to the function of BLA in mice can be understood to apply to primates. Importantly, the patterns of projections from BLA to the nucleus accumbens were prominent in both animals (**Figure 1D**), indicating a somewhat conserved functional circuit. By contrast, the organization of single BLA neuron projections to frontal cortex differed markedly between mice and macaques, and in some cases these differences matched known functional distinctions between species in medial frontal cortex^19^. In other cases, however, ventral frontal areas received quite different inputs from BLA despite their highly similar functions^27^. These results highlight that appropriate interpretation of cross-species behavioral neuroscience findings requires nuance and provide a basis for future comparative studies.

## Online methods

### Subjects

Two male rhesus macaques (*Macaca mulatta*) 8-9 years of age, were used for our experiments, weighing between 10 and 15 kg; these are the same animals reported in ref.^16^. Animals were housed individually and kept on a 12-hour light/dark cycle. Food was provided daily with water *ad libitum*. Environmental enrichment was provided daily, in the form of play objects or small food items. Five laboratory mice (*Mus musculus,* C57Bl/6J) aged P42-P60 were acquired from Jackson Laboratories and used for these experiments. All procedures were approved by the Icahn School of Medicine IACUC. Procedures involving macaques were carried out in accordance with NIH standards for work involving non-human primates.

### Virus preparation

Modified sindbis virus for MAPseq was obtained from the MAPseq Core Facility at Cold Spring Harbor Laboratory (CSHL)^15^. The viral library used in this study had a diversity of 20,000,000 unique barcodes. All viruses were stored at -80°C and aliquots were thawed over wet ice immediately prior to injection.

### Surgery and perfusion

#### Macaques

The surgical approach is detailed in Zeisler et al.^16^. Briefly, for each animal, T1-weighted MRIs were collected to localize the basal and lateral nuclei of the amygdala. Twelve to 15 injections of 0.4 μl were delivered at a rate of 0.2 μl per minute using Hamilton syringes, and animals were perfused 67-72 hours after completing injections with 1% formaldehyde (from paraformaldehyde, PFA; Electron Microscopy Sciences) in phosphate-buffered saline (PBS, Invitrogen) for two minutes followed by 4% PFA for 18 minutes.

#### Mice

Subjects were anesthetized with isoflurane and mounted in a small animal stereotaxic apparatus. Two injections of sindbis vector (0.15 μl each) were made in each hemisphere using a Neurosyringe and Stoelting motorized injector through a small craniotomy, relying on coordinates that have been empirically tested within the Clem laboratory^39–41^. The scalp incision was closed with sutures and VetBond tissue adhesive. Banamice (2.5 mg/kg) was administered subcutaneously for postoperative analgesia. Perfusion took place 40-44 hours after viral injections. Mice were deeply anesthetized via isoflurane inhalation and transcardially perfused with 35-40 ml 4% formaldehyde (diluted fresh from 20% PFA; Electron Microscopy Sciences) over 15-20 minutes.

#### Tissue preparation

As previously described^16^, macaque brains were blocked by separating the hemispheres, removing the temporal lobe, and making a single coronal cut caudal to the thalamus. Blocks were post-fixed in 4% PFA in PBS at 4°C then were frozen on dry ice and stored at -80°C before sectioning. For mice, brains were post-fixed for 24 hours in 4% PFA at 4°C shaken continuously, after which they were rinsed with PBS, dried, frozen on dry ice, and stored at -80°C before sectioning.

### Sectioning and dissection

For both mice and macaque tissue, brain blocks were sectioned at 200 μm on a Leica 3050S cryostat over dry ice and stored at -80°C prior to dissection. Cortical areas were then dissected according to anatomical landmarks over dry ice. The areas that were collected in macaques and our operational definitions based on sulci can be found in **Table 1**. In mice, we took tissue punches from areas based on the 2^nd^ edition of the Paxinos and Franklin atlas^42^ using anterior-posterior and medio-lateral anatomical markers to guide punches; see **Table 2** for detailed locations of punches.

In macaques, samples from each area were combined across 3 sections in the anterior-posterior plane into 1.5 ml Eppendorf tubes, which were stored at -80°C prior to shipping frozen on dry ice for sequencing.

In mice, dissected punches were combined across areas into 1.5 ml Eppendorf tubes (such that all punches from a single area were collected in the same tube) and stored at -80°C prior to shipping frozen on dry ice for sequencing.

### mRNA sequencing and preprocessing

Sequencing of MAPseq projections was performed by the MAPseq Core Facility at CSHL as described in Kebschull et al.^15^. Briefly, RNA was extracted using a Trizol-based protocol, RNA quality verified by Bioanalyzer, and qPCR performed to assess barcode quantity. Then, double-stranded cDNA was synthesized and submitted for sequencing using the Illumina NextSeq platform. Preprocessing by CSHL results in a barcode matrix with size n samples x n barcodes.

### Filtering and analysis

Filtering was performed in Python 3.9^43–46^ as described previously^16^; specific analyses can be found in the code available on Github. Briefly, barcodes survive filtering if the max counts recovered from the injection site is greater than 20 and the max counts from at least one target site is greater than 5^16^. Then, barcode matrices were binarized and collapsed within brain regions (e.g., if one of the eight entorhinal cortex samples had enough barcode, then we interpret that neuron as projecting to the entorhinal cortex). Finally, within each species, animal-specific barcode matrices were combined into a single population for future analyses.

Projection strength plots (Figure 1B) were computed by summing the number of unique barcodes found in each brain region and dividing by the total number of barcodes recovered in each species. The number of targets for each neuron (Figure 1C) was computed by counting the number of distinct brain regions in which sufficient barcode was recovered. The conditional probabilities (Figure 1D) were calculated by finding the subset of cells in each area which project to each of the remaining areas. Thus, the values plotted indicate the probability that a neuron found in some area A is also found in another area B: P(B|A).

In the network analysis figures (Figure 2 and Supplemental Figure 2), we filtered for neurons which projected to medial frontal cortex (scACC, pgACC, and dACC in the macaque; IL, PL, CG1/2 in the mouse) or ventral frontal cortex (mOFC, lOFC, vlPFC, and AI in the macaque; MO, LO, VO, and AI in the mouse). While projections to CG1 and CG2 were analyzed separately previously, to compare to dACC more accurately^47^, neurons projecting to CG1 and CG2 were combined into a single population by including those which projected to one area or both. First, we computed the degree of overlap within these networks by preparing Venn diagrams of projections specific to these areas and how many neurons branched between them. The proportion of branching within each area was compared using z-tests for proportions; p-values were adjusted using FDR correction. We then focused on the projections which were specific by excluding any neurons which projected to multiple areas in the same network (i.e. the areas on the outside of the Venn diagram). Projection strength to other areas were calculated as described above and compared using pairwise z-tests for proportions, corrected for multiple comparisons.

Finally, in the control analyses in Supplemental Figure 1, we first counted the proportion of recovered barcodes which were found in control sites in cerebellum or primary visual cortex in macaques and mice, respectively. Then, as we had done previously^16^, we assessed the effects of filtering parameters on recovery of mouse barcodes by iteratively re-filtering with different thresholds. Finally, using *np.unique*, we assessed the frequency of unique projection patterns as a proxy for the proportion of neurons which were infected with a single unique barcode sequence.

## AUTHORS CONTRIBUTIONS

Conceptualization and Methodology: ZZ, KAH, FMS, PRH, RLC, PHR; Investigation, Data curation, Analysis, Software and Visualization: ZZ, KAH, RLC, PHR; Writing – Original Draft and Review/Editing: ZZ, KAH, FMS, PRH, RLC, PHR; Funding Acquisition and Supervision: RLC and PHR.

## ACKNOWLEDGEMENTS

This work was supported by a National Institute of Neurological Disorders and Stroke award to RLC and PHR (R34NS122050) and seed funds from the Icahn School of Medicine at Mount Sinai to PHR. We would like to thank Alex Vaughan and Tony Zador for advice and encouragement, and we are grateful to Steven P Wise and members of the Rudebeck and Clem laboratories for discussion and comments on earlier versions of the manuscript.

## References

1. Bucy, P. C. & Klüver, H. An anatomical investigation of the temporal lobe in the monkey (Macaca mulatta). Journal of Comparative Neurology 103, 151–251 (1955).

2. Horvath, F. E. Effects of basolateral amygdalectomy on three types of avoidance behavior in cats. Journal of Comparative and Physiological Psychology 56, 380–389 (1963).

3. Sengupta, A. et al. Basolateral Amygdala Neurons Maintain Aversive Emotional Salience. J. Neurosci. 38, 3001–3012 (2018).

4. Schumann, C. M., Bauman, M. D. & Amaral, D. G. Abnormal structure or function of the amygdala is a common component of neurodevelopmental disorders. Neuropsychologia 49, 745–759 (2011).

5. Brady, R. O. et al. State dependent cortico-amygdala circuit dysfunction in bipolar disorder. Journal of Affective Disorders 201, 79–87 (2016).

6. Ghashghaei, H. T., Hilgetag, C. C. & Barbas, H. Sequence of information processing for emotions based on the anatomic dialogue between prefrontal cortex and amygdala. NeuroImage 34, 905–923 (2007).

7. Hintiryan, H. et al. Connectivity characterization of the mouse basolateral amygdalar complex. Nat Commun 12, 2859 (2021).

8. Price, J. L. & Amaral, D. G. An autoradiographic study of the projections of the central nucleus of the monkey amygdala. J. Neurosci. 1, 1242–1259 (1981).

9. Amaral, D. G. & Price, J. L. Amygdalo-cortical projections in the monkey (Macaca fascicularis). Journal of Comparative Neurology 230, 465–496 (1984).

10. Janak, P. H. & Tye, K. M. From circuits to behaviour in the amygdala. Nature 517, 284–292 (2015).

11. Chareyron, L. J., Lavenex, P. B., Amaral, D. G. & Lavenex, P. Stereological analysis of the rat and monkey amygdala. Journal of Comparative Neurology 519, 3218–3239 (2011).

12. Wise, S. P. Forward frontal fields: phylogeny and fundamental function. Trends in Neurosciences 31, 599–608 (2008).

13. Murray, E. A., Wise, S. P. & Graham, K. S. The evolution of memory systems: ancestors, anatomy, and adaptations. (Oxford university press, 2017).

14. Passingham, R. E. & Wise, S. P. The neurobiology of the prefrontal cortex: anatomy, evolution, and the origin of insight. (Oxford University Press, 2012).

15. Kebschull, J. M. et al. High-Throughput Mapping of Single-Neuron Projections by Sequencing of Barcoded RNA. Neuron 91, 975–987 (2016).

16. Zeisler, Z. R. et al. Single basolateral amygdala neurons in macaques exhibit distinct connectional motifs with frontal cortex. Neuron 111, 3307–3320.e5 (2023).

17. Gergues, M. M. et al. Circuit and molecular architecture of a ventral hippocampal network. Nature Neuroscience 1–9 (2020) doi:10.1038/s41593-020-0705-8.

18. Han, Y. et al. The logic of single-cell projections from visual cortex. Nature 556, 51–56 (2018).

19. Wallis, C. U., Cardinal, R. N., Alexander, L., Roberts, A. C. & Clarke, H. F. Opposing roles of primate areas 25 and 32 and their putative rodent homologs in the regulation of negative emotion. PNAS 114, E4075–E4084 (2017).

20. Alexander, L., Wood, C. M. & Roberts, A. C. The ventromedial prefrontal cortex and emotion regulation: lost in translation? The Journal of Physiology **n/a**,.

21. Dal Monte, O., Chu, C. C. J., Fagan, N. A. & Chang, S. W. C. Specialized medial prefrontal– amygdala coordination in other-regarding decision preference. Nature Neuroscience 1–10 (2020) doi:10.1038/s41593-020-0593-y.

22. Dal Monte, O., et al. Widespread implementations of interactive social gaze neurons in the primate prefrontal-amygdala networks. Neuron 110, 2183–2197.e7 (2022).

23. Smith, M. L., Asada, N. & Malenka, R. C. Anterior cingulate inputs to nucleus accumbens control the social transfer of pain and analgesia. Science 371, 153–159 (2021).

24. Schneider, K. N., Sciarillo, X. A., Nudelman, J. L., Cheer, J. F. & Roesch, M. R. Anterior Cingulate Cortex Signals Attention in a Social Paradigm that Manipulates Reward and Shock. Current Biology 30, 3724–3735.e2 (2020).

25. Arruda-Carvalho, M. & Clem, R. L. Pathway-Selective Adjustment of Prefrontal-Amygdala Transmission during Fear Encoding. J. Neurosci. 34, 15601–15609 (2014).

26. Alexander, L. et al. Over-activation of primate subgenual cingulate cortex enhances the cardiovascular, behavioral and neural responses to threat. Nature Communications 11, 5386 (2020).

27. Rudebeck, P. H. & Izquierdo, A. Foraging with the frontal cortex: A cross-species evaluation of reward-guided behavior. Neuropsychopharmacol. 47, 134–146 (2022).

28. Heilbronner, S. R., Rodriguez-Romaguera, J., Quirk, G. J., Groenewegen, H. J. & Haber, S. N. Circuit-Based Corticostriatal Homologies Between Rat and Primate. Biological Psychiatry 80, 509–521 (2016).

29. Walter, N. et al. Effect of chronic ethanol consumption in rhesus macaques on the nucleus accumbens core transcriptome. Addiction Biology 26, e13021 (2021).

30. Stacey, D. et al. A translational systems biology approach in both animals and humans identifies a functionally related module of accumbal genes involved in the regulation of reward processing and binge drinking in males. Journal of Psychiatry and Neuroscience 41, 192–202 (2016).

31. Bracht, T. et al. The role of the orbitofrontal cortex and the nucleus accumbens for craving in alcohol use disorder. Transl Psychiatry 11, 1–10 (2021).

32. Taswell, C. A., Costa, V. D., Murray, E. A. & Averbeck, B. B. Ventral striatum’s role in learning from gains and losses. Proceedings of the National Academy of Sciences 115, E12398–E12406 (2018).

33. Fraser, K. M. & Janak, P. H. Long-lasting contribution of dopamine in the nucleus accumbens core, but not dorsal lateral striatum, to sign-tracking. European Journal of Neuroscience 46, 2047– 2055 (2017).

34. Zimmermann, K. S., Yamin, J. A., Rainnie, D. G., Ressler, K. J. & Gourley, S. L. Connections of the Mouse Orbitofrontal Cortex and Regulation of Goal-Directed Action Selection by Brain-Derived Neurotrophic Factor. Biol Psychiatry 81, 366–377 (2017).

35. Winstanley, C. A., Theobald, D. E. H., Cardinal, R. N. & Robbins, T. W. Contrasting Roles of Basolateral Amygdala and Orbitofrontal Cortex in Impulsive Choice. J Neurosci 24, 4718–4722 (2004).

36. Aguirre, C. G. et al. Dissociable contributions of basolateral amygdala and ventrolateral orbitofrontal cortex to flexible learning under uncertainty. J. Neurosci. (2023) doi:10.1523/JNEUROSCI.0622-23.2023.

37. Rudebeck, P. H., Saunders, R. C., Lundgren, D. A. & Murray, E. A. Specialized Representations of Value in the Orbital and Ventrolateral Prefrontal Cortex: Desirability versus Availability of Outcomes. Neuron 95, 1208–1220.e5 (2017).

38. Price, J. L. Definition of the Orbital Cortex in Relation to Specific Connections with Limbic and Visceral Structures and Other Cortical Regions. Annals of the New York Academy of Sciences 1121, 54–71 (2007).

39. Cummings, K. A. & Clem, R. L. Prefrontal somatostatin interneurons encode fear memory. Nat Neurosci 23, 61–74 (2020).

40. Lucas, E. K., Jegarl, A. M., Morishita, H. & Clem, R. L. Multimodal and Site-Specific Plasticity of Amygdala Parvalbumin Interneurons after Fear Learning. Neuron 91, 629–643 (2016).

41. Lucas, E. K., Wu, W.-C., Roman-Ortiz, C. & Clem, R. L. Prazosin during fear conditioning facilitates subsequent extinction in male C57Bl/6N mice. Psychopharmacology 236, 273–279 (2019).

42. Paxinos, G., Franklin, K. B. J. & Franklin, K. B. J. The mouse brain in stereotaxic coordinates. (Academic Press, 2001).

43. Harris, C. R. et al. Array programming with NumPy. Nature 585, 357–362 (2020).

44. Pedregosa, F. et al. Scikit-learn: Machine Learning in Python. Journal of Machine Learning Research 12, 2825–2830 (2011).

45. Hunter, J. D. Matplotlib: A 2D Graphics Environment. Computing in Science & Engineering 9, 90– 95 (2007).

46. Waskom, M. L. seaborn: statistical data visualization. Journal of Open Source Software 6, 3021 (2021).

47. van Heukelum, S. et al. Where is Cingulate Cortex? A Cross-Species View. Trends in Neurosciences 43, 285–299 (2020).

