## Supplemental figures and legends for "Comparative basolateral amygdala connectomics reveals dissociable single-neuron projection patterns to frontal cortex in macaques and mice"

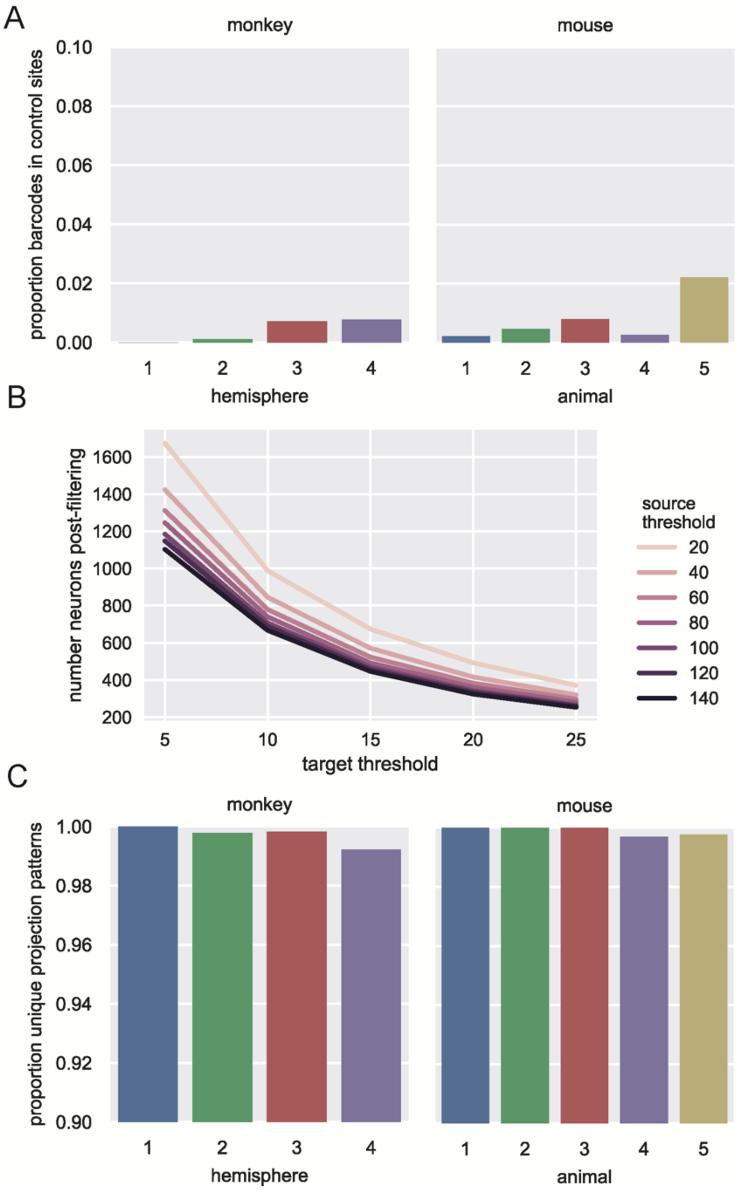


**Supplemental Figure 1. Control analyses confirm similar efficacy of MAPseq in both mice and macaques.** A) Few barcodes were recovered from control sites in cerebellum (macaques; left) and primary visual cortex (mice; right), indicating the specificity of MAPseq barcode infection and transport. B) Filtering thresholds have little effect on number of recovered barcodes in mice. C) Barcode expression patterns were primarily unique, suggesting that infection of single neurons by more than one barcode is exceedingly rare in both macaques (left) and mice (right).

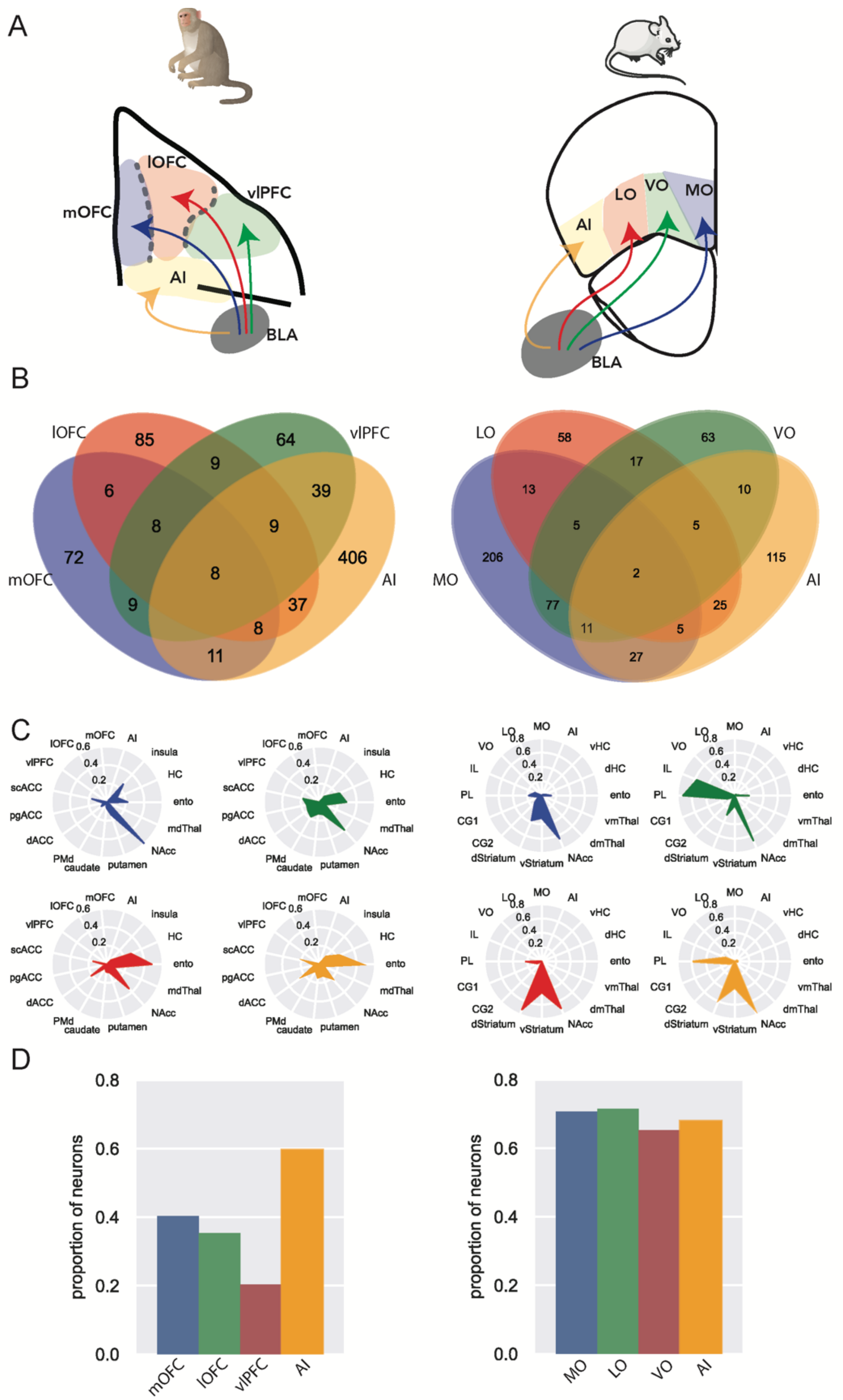


**Supplemental Figure 2.** A) Schematic of populations of ventral-projecting neurons in macaques (left; ventral view) and mice (right; coronal view). B) Venn diagrams of within-ventral frontal cortex branching. Macaque AI received the most specific BLA input (AI vs mOFC: z = 4.43, $p < 0.0001$), while MO- and AI- projecting neurons in the mouse branched least within ventral frontal cortex (AI vs LO: z=

2.29, $p = 0.220$; MO vs LO: z = 2.919, $p = 0.004$). vlPFC- and VO-projecting neurons were most likely to branch within ventral frontal cortex (vlPFC vs mOFC: z = 2.48, $p = 0.013$; VO vs LO: z = 2.076, $p = 0.379$). C) Likelihood of ventral frontal-projecting neurons branching to project to other areas. This plot excludes those neurons which branch between multiple ventral frontal areas, such that the populations are non-overlapping. LO-projecting neurons were highly likely to branch to PL (LO vs VO: z.= 4.053, $p < 0.0001$) and IL (LO vs VO: z = 2.507, $p = 0.012$), while lOFC-projecting neurons were least likely to also project to pgACC (lOFC->pgACC vs lOFC->scACC: z = 3.642, $p = 0.0003$). D) Likelihood of ventral frontal-projecting neurons projecting to NAcc. While macaque AI-projecting neurons were more likely than all other ventral areas to also project to NAcc (Fisher's exact test: AI vs mOFC, $p = 0.0057$; AI vs lOFC, $p = 0.00019$; AI vs vlPFC, $p < 0.0001$), all mouse ventral frontal areas were equally and highly likely to project to NAcc (LO vs VO: $p = 1.0$).
